# The genetic architecture of milk urea concentration in dairy cattle differs across the lactation cycle

**DOI:** 10.64898/2026.04.22.719978

**Authors:** Qiongyu He, Stefan Vasiljevic, Naveen Kumar Kadri, Natasha Watson, Patrick Stratz, Xena Marie Mapel, Alexander S. Leonard, Franz R. Seefried, Hubert Pausch

## Abstract

Milk urea concentration (MUC) is an indicator of dietary protein utilization and nitrogen use efficiency in dairy cows. We performed genome-wide association studies (GWAS) on MUC in early, mid, and late lactation in the Holstein (HOL) and Brown Swiss (BSW) dairy cattle breeds using imputed sequence variants. We identified 11 and 17 independent quantitative trait loci (QTL) for MUC across the three lactation stages in BSW and HOL, respectively. While many of these QTL have previously been reported for MUC and other dairy traits, our study provides evidence that some QTL exert lactation-stage specific effects. Our findings suggest that variants at the *DGAT1* locus on BTA14 have pleiotropic effects on MUC and other dairy traits. This QTL showed an early lactation-specific association with MUC but impacted milk and fat yield across the entire lactation. We fine-mapped two QTL for MUC in early and mid-lactation in BSW on BTA9 (lead SNP: 9:21392941, P_corrected_ = 1.1E-17) and BTA28 (lead SNP: 28:6518357; P_corrected_ = 3E-11). We identified lncRNA ENSBTAG00000058688 and *IBTK* as positional and functional candidate genes for the BTA9 QTL, and *KCNK1* as positional and functional candidate gene that harbors a highly significant missense variant for the BTA28 QTL. In conclusion, our results shed light on the genetic architecture of MUC and highlighted QTL harboring potential functional variants underpinning milk urea variation within and across breeds.

## Introduction

Milk urea concentration (MUC) is a reliable indicator of nitrogen use efficiency and nitrogen excretion in dairy cattle (Powell et al., 2011; Barros et al., 2019). MUC is proportional to blood urea concentration, which predominantly derives from systemic protein metabolism (Butler et al., 1996; Ghavi Hossein-Zadeh, 2024). Therefore, MUC reflects the activity of multiple biological pathways involved in protein synthesis, nitrogen metabolism, and the animal’s energy balance (Oltner and Wiktorsson, 1983; Westwood et al., 1998; Ghavi Hossein-Zadeh, 2024). MUC is likely to share part of its genetic architecture with other milk composition traits, since pathways of fat metabolism and protein synthesis jointly regulate nitrogen utilization and milk composition (Hojman et al., 2004; Rzewuska and Strabel, 2013; Jahnel et al., 2023).

Previous studies have demonstrated that MUC is moderately heritable (Buitenhuis and Poulsen, 2023). Selection for lower MUC may reduce urinary nitrogen excretion and improve environmental sustainability (O’Callaghan et al., 2019; Marshall et al., 2021; Prahl et al., 2022). However, too low MUC can reflect inadequate protein status and impaired nitrogen metabolism, potentially compromising health and reproductive performance (Hojman et al., 2004). Therefore, maintaining an optimal MUC is essential for balancing productivity and health with excess nitrogen excretion (Zhao et al., 2025).

Several genome-wide association studies (GWAS) have reported quantitative trait loci (QTL) for MUC in cattle (Pegolo et al., 2018; Ma et al., 2023; Atashi et al., 2024). Multi-breed GWAS (Van Den Berg et al., 2022; Atashi et al., 2024) identified MUC-associated variants show consistent effects across breeds, suggesting a shared genetic architecture. These studies for MUC relied on global phenotypes that were estimated over the entire lactation, which precluded the investigation of stage-specific genetic effects underlying dynamic changes in MUC across lactation, including those related to metabolic, endocrine, and immune regulation (Bionaz and Loor, 2008; Mortazavi et al., 2025). While the lactation cycle has pronounced effects on milk yield and composition (Lund et al., 2008; Strucken et al., 2012), lactation stage-specific analyses for MUC remain relatively underexplored (Mortazavi et al., 2025). Using MUC measured at distinct lactation stages may help resolve the temporal genetic architecture of dairy traits across the lactation cycle.

Here, we investigated the genetic architecture of MUC in early, mid, and late lactation in two dairy breeds. We identified previously reported and novel QTL including several with lactation stage-specific effects. We further identified positional and functional candidate genes for two QTL on BTA9 and BTA28 and prioritized a missense variant in *KCNK1* as a candidate causal variant for MUC in cattle.

## Materials and Methods

### Phenotype

Raw phenotypes for milk urea concentration (MUC; mg/dL) were derived from mid-infrared (MIR) spectral (International Committee for Animal Recording (ICAR), 2020) predictions of milk samples collected across multiple breeds in accordance with ICAR guidelines (International

Committee for Animal Recording (ICAR), 2023). The raw data were provided by Swissherdbook, Holstein Switzerland, and Braunvieh Schweiz. Test-day records were available across all parities. After excluding records with MUC < 5 mg/dL, 31,259,203 test-day records remained for the Braunvieh group, comprising Brown Swiss (BSW) and Original Braunvieh (OB), whereas 48,791,193 remained for the Holstein group, comprising Holstein (HOL), Swiss Fleckvieh (SF), and Simmental (SIM), for further processing. These data were subsequently used to estimate variance components using a multi-trait random regression test-day animal model (RRTDM) (Jamrozik and Schaeffer, 1997), processed by Qualitas AG (Zug, Switzerland), thereby enabling the genetic evaluation of MUC as part of the Swiss routine genetic evaluation system.

In the RRTDM, MUC was jointly analyzed with the routinely evaluated dairy traits milk yield (MKG; kg), fat yield (FKG; kg), and protein yield (PKG; kg) in the Braunvieh and Holstein groups. The model accounts for systematic environmental effects and lactation-stage-dependent changes in test-day performance across days in milk (DIM). Specifically, herd-test-day (HTD) and time-region-age-parity-season (TRAPS) (Stratz et al., 2025) were fitted as fixed effects, with TRAPS modeled over DIM using orthogonal Legendre polynomials of order 0 to 6 (Schaeffer, 2016). Additive genetic and permanent environmental effects were modeled by random regression using Legendre polynomials up to order 4. Residual variance was assumed heterogeneous across 4 DIM classes within parity: 5–45, 46–115, 116–265, and 266–365 d. Parity 1 to 3 were modeled explicitly, and later parities were treated as repeated records of parity 3.

Model parameters were estimated in a Bayesian framework using Gibbs sampling (Jamrozik et al., 1997). To assess the stability of the estimates, the filtered test-day dataset was divided into multiple randomly sampled datasets at the herd-test-day level. For each sampled dataset, 100,000

Gibbs sampling iterations were run to obtain posterior distributions of additive genetic, permanent environmental, and residual (co)variance components, and the corresponding posterior means were retained for downstream calculations. Burn-in was determined on one randomly selected sampled dataset using the Heidelberger-Welch convergence diagnostic implemented in the R package coda (Plummer et al., 2006), evaluated every 100th iteration, and the resulting burn-in was then applied to all sampled datasets. The resulting estimates were used to derive daily heritability of MUC across DIM (Supplemental Figure S1; see Notes) for routine evaluation. For this study, yield deviations (YD), defined as test-day records adjusted for the fitted HTD and TRAPS effects from RRTDM, were used as input phenotypes for the subsequent genome-wide association study (GWAS). The YD were defined as:

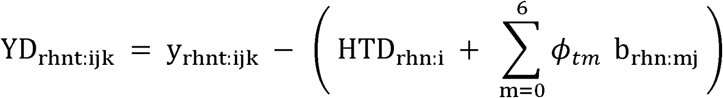

where r, h, n, t, i, j, and k are indices for breed (Braunvieh or Holstein group), trait, parities (1,2, or ≥ 3), DIM, HTD subclass, TRAPS subclass, and cow, respectively. HTD_rhn:i_ is the fitted HTD effect, b_rhn:mj_ are the fitted TRAPS regression coefficients, and *ϕ_tm_* is the orthogonal Legendre polynomial covariate of order m _(1..6)_ evaluated at DIM t.

For the first parity, we obtained 16,218,745 YD records from 2,152,381 cows, of which records for BSW and HOL cows were retained for downstream GWAS. These records for MUC were divided into ten consecutive 30-day intervals, starting from day 10. Cows with at least one adjusted record within a given interval were retained. To obtain a single record per animal, observations were averaged within each interval. We selected windows 1, 5 and 10 for further exploration of the genetic architecture of MUC, representing early, mid, and late lactation. The number of observations was higher for window 1 (BSW: 23,243; HOL: 17,736) than window 5 (BSW: 19,715; HOL: 15,306) and window 10 (BSW: 13,729; HOL: 10,980). The same approach was used to obtain phenotypes for FKG, PKG, and MKG.

### Genotypes

We considered 178,845 animals from the BSW, OB, HOL, SIM and SF breeds that had microarray-derived genotypes. The animals were genotyped using 10 arrays comprising between 20K and 777K SNPs. Quality control was performed for each array type separately as described in Watson et al (Watson et al., 2025). Subsequently, genotype imputation was performed in a stepwise procedure with Beagle (v 5.4) (Browning et al., 2021) for two breed groups separately: BSW, OB, and their crosses, and HOL, SIM, SF, and their crosses. In the first step, Illumina Bovine HD SNP BeadChip genotypes were imputed into low- and medium-density array-derived genotypes using reference panels of 1,183 BSW/OB and 1,138 HOL/SIM/SF animals. In the second step, whole-genome sequence data from 607 in-house sequenced BSW and OB (Lloret-Villas et al., 2023), and 1,464 HOL and SIM animals selected from the 1000 Bull Genomes Project run 9 (Daetwyler et al., 2014) were used as a reference to impute sequence-level variants. This approach provided genotypes for 26,265,006 sequence variants for the BSW and OB group, and 37,153,418 sequence variants for the HOL, SIM, and SF group. For the subsequent analyses, we retained 60,564 BSW and 89,315 HOL animals.

### Genetic parameter estimation on genomic data

We built genomic relationship matrices (GRMs) using the “--grm” flag of GCTA (v1.94.1) (Yang et al., 2011) considering genotypes at 14,576,969 and 14,094,138 sequence variants in BSW and HOL, respectively, that had minor allele frequency (MAF) greater than 0.5% and imputation accuracy (Beagle’s r^2^) greater than 0.4. The principal components (PC) of the GRM were obtained with the “-pca” flag of GCTA. Genome-wide additive heritability (h^2^) for the window-specific traits was estimated with the REML algorithm (“-reml”-flag) of GCTA, and genetic correlations between windows were estimated with the “--reml-bivar" option.

### Genome-wide association study

Additive GWAS were conducted between the imputed sequence variant genotypes and the window-specific traits within the two breeds separately using a mixed linear model-based approach implemented in the BOLT-LMM (v2.4.1) software (Loh et al., 2015). We retained SNPs that had MAF ≥ 0.01 in each lactation stage-specific cohort. Linkage disequilibrium (LD) scores were computed with PLINK (v1.9) (Purcell et al., 2007). The LD-pruned subset considered to construct the model SNP (Loh et al., 2018) contained 1,460,499 and 1,686,281 SNPs in BSW and HOL, respectively. The top four PCs were included as covariates for association testing. SNPs that deviated from Hardy-Weinberg proportions (P <E-05) were removed. Residual inflation detected in the GWAS summary statistics was accounted for using genomic control (Devlin and Roeder, 1999).

Association analysis conditional on the most significantly associated marker was performed within breeds using the “-cojo” flag of GCTA (Genetic Investigation of ANthropometric Traits (GIANT) Consortium et al., 2012). Mendelian randomization (MR) was performed with the GSMR (Zhu et al., 2018) model of GCTA to investigate a putative causal association between MKG and MUC or FKG and MUC. For each trait pair, both forward and reverse GSMR analyses were conducted (--gsmr-direction 2). LD-clumped (r² = 0.5) genome-wide significant SNPs (P_GWAS_ < 5E-08) were used as instrumental variables (IVs), and the HEIDI outlier test (--heidi-thresh 0.1) was applied to remove variants whose effects were inconsistent with the main MR signal. To account for multiple testing, the significance threshold was defined as P_GSMR_ < 0.05 / N_IV_ (Schwarz et al., 2025). Variants within ±2 Mb of the lead SNP (14:598800) at the BTA14 QTL (0–2,598,800 bp) were excluded to investigate the contribution of this locus to the GSMR-derived causal association between MUC and MKG.

### Functional annotation

Sequence variants were annotated using the Ensembl Variant Effect Predictor (McLaren et al., 2016) based on the bovine reference genome (ARS-UCD1.2) and the corresponding Ensembl annotation (Dyer et al., 2025). The pairwise LD between the lead SNPs and all other SNPs within ±1 Mb was calculated in the full cohort of animals from both breeds using PLINK. We used FIMO from MEMESuite (Bailey et al., 2009) to identify putative transcription factor (TF) binding motifs from the Cis-BP database (Weirauch et al., 2014). We selected the most significant predicted TF binding site for each genomic region. Only sites with a Bonferroni-corrected p-value less than 0.05 were retained. The GC-content was calculated based on the respective chromosomes.

## Results and Discussion

### Milk urea concentration across the lactation cycle

Across the entire lactation cycle, MUC was slightly higher in BSW than in HOL, averaging 26.39 ± 8.79 mg/dL and 23.80 ± 8.57 mg/dL, respectively. MUC was subsequently assessed in early, mid, and late lactation in BSW and HOL cattle using test-day records adjusted for the fixed effects of HTD and TRAPS. In both breeds, MUC yield deviations were the lowest in early lactation, whereas the corresponding standard deviations were relatively similar across stages (Supplemental Table S1; see Notes).

The genomic heritability (h²) of MUC was moderate in both breeds (Table 1), which aligns with previous studies in dairy cattle (Buitenhuis and Poulsen, 2023). In both breeds, h^2^ fluctuated mildly across the lactation cycle and was lowest in early lactation and highest in mid lactation. The genetic correlation of MUC between the lactation stages was high (r = 0.70 - 0.97; Supplemental Figure S2; see Notes) and strongest between mid and late lactation. The genetic correlation between MUC and other dairy traits (MKG, FKG, PKG) was low in both breeds (Supplemental Figure S2; see Notes).

**Table 1:** Genomic heritability of milk urea concentration in 2 cattle breeds.

| Breeds | Early lactation (DIM 10-40) | Mid lactation (DIM 130-160) | Late lactation (DIM 280 - 310) |
| --- | --- | --- | --- |
| BSW | $0.1597 \pm 0.0083$<br>(n=23,243) | $0.2004 \pm 0.0096$<br>(n=19,715) | $0.1872 \pm 0.0114$<br>(n=13,729) |
| HOL | $0.1534 \pm 0.0092$<br>(n=17,736) | $0.1883 \pm 0.0107$<br>(n=15,306) | $0.1829 \pm 0.0127$<br>(n=10,980) |

Additive heritability and corresponding standard errors ( $h^2 \pm SE$ ) of MUC in early (DIM 10–40), mid (DIM 130–160), and late (DIM 280–310) lactation in BSW and HOL cattle. Sample sizes (n) represent the number of animals with test-day records that were used to estimate $h^2$ .

### GWAS reveal lactation cycle-specific QTL for milk urea concentration

In BSW, our lactation stage-specific GWAS revealed 4, 6 and 1 independent QTL (P_corrected_ < 5E-08, P_cojo_ < 5E-08) for MUC in early, mid and late lactation, respectively. Correspondingly, 6, 6, and 5 independent QTL were identified in HOL (Figure 1) (Supplemental Table S2; see Notes). While several of the MUC QTL overlapped genomic regions associated with milk yield, fat yield, or protein yield (Supplemental Figure S3; see Notes), the overwhelming majority of the MUC QTL lead variants were not associated with any of the three dairy traits. Differences in sample size and MUC h^2^ were observed across the three lactation stages. The statistical power to detect QTL for MUC was similar for the early and mid-lactation in both breeds, but was reduced in late lactation (Wu et al., 2022), which likely limited the detection of significant QTL during this stage.

**Figure 1:**
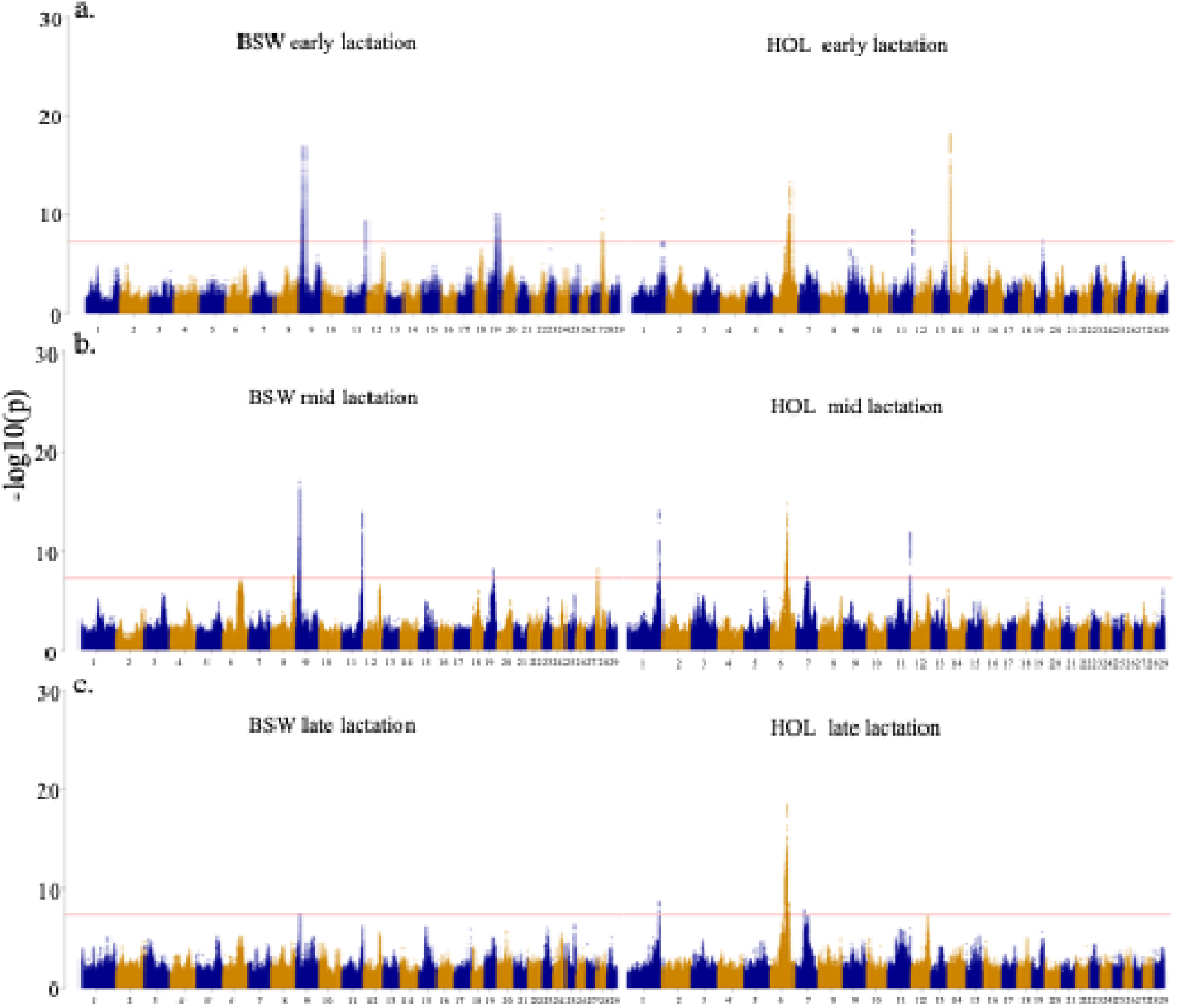
Results of lactation stage-specific GWAS for MUC in BSW and HOL cattle. Manhattan plots represent the association of imputed sequence variants with milk urea concentration, in early (a), mid (b) and late (c) lactation in BSW (left) and HOL (right). The red line shows the significance threshold of −log10 (5E-08).

Several QTL for MUC overlapped genomic regions that had previously been reported to be associated with MUC and other dairy traits. However, unlike earlier studies that relied on global phenotypes averaged across the entire lactation cycle, our GWAS used lactation stage–specific phenotypes, thereby providing a more nuanced view of how these regions might influence MUC across the lactation cycle. This included a QTL in BSW (BTA19, ∼43.9 Mb) (Cecchinato et al., 2014; Lopdell, 2023), and two QTL in HOL (BTA14, ∼0.6 Mb; BTA19, 41.7Mb) (Cecchinato et al., 2014; Van Den Berg et al., 2022; Lopdell, 2023; Ma et al., 2023; Atashi et al., 2024), which primarily affected MUC in the early lactation. Additionally, a BSW QTL on BTA8 (103.4 Mb) (Chen et al., 2023) primary affected MUC in mid lactation.

We also identified QTL that were significant across multiple lactation stages. Two HOL QTL on BTA1 (146.8 Mb) (Atashi et al., 2022) and BTA6 (∼85.6 Mb) (Pegolo et al., 2018; Atashi et al., 2024) were detected across all three stages while a QTL on BTA11 (103.2 Mb) (Atashi et al., 2024) was observed in both BSW and HOL in early and mid-lactation only. For all QTL that were shared across lactation stages, the lead variant from a specific window was typically also highly associated with other windows, albeit it was not necessarily the most significant variant. Variation in statistical power across the three lactation stages may partially explain the difference in significance for some QTL; specifically, the stronger signal for BTA1 and BTA11 at the mid stage than the early stage, and the weaker late-lactation signal at the BTA11 QTL. However, a stronger late-than-early signal at the BTA1 QTL in HOL cannot be explained with differences in statistical power and likely indicates late lactation stage-specificity. The lead variants for the mid (1:146904166) and late (1:146847984) lactation QTL (Supplemental Table S2; see Notes) reside nearby *LOC132346140* encoding chloride intracellular channel protein 6-like and overlap a milk urea nitrogen QTL identified in an earlier study (Van Den Berg et al., 2022).

### A QTL on BTA14 in HOL has potential horizontal pleiotropic effects

The GWAS for MUC identified a strong QTL for early lactation in HOL at the proximal region of BTA14 (lead SNP: 14:598800, rs431919726, P_corrected_=5.5E-19;). The rs431919726 variant is close to the K232A-variant of *DGAT1* (Van Den Berg et al., 2016), which has a large impact on milk yield and composition (Winter et al., 2002; Pausch et al., 2017). The signal was absent when the association analysis was conditioned on the causal *DGAT1* polymorphism 14:611019 (r2=0.97) and 14:611020 (r2=0.97), suggesting that the K232A-variant also affects MUC. The milk yield–decreasing but fat yield–increasing lysine variant of the *DGAT1* p.K232A polymorphism increased MUC. While the association of this QTL with MUC was primarily apparent in the early lactation, it was significant for FKG and MKG across all lactation stages (Supplemental Figure S3; see Notes) (Liu et al., 2022a), possibly suggesting that distinct regulatory pathways affect MUC, MKG and FKG.

The rs431919726-variant was also in high LD (r² = 0.97) with the lead variant (14:544162) at a *DGAT1* liver eQTL (P_eQTL_ = 5.97 E-16) and the lead variant (14:568472) at a *DGAT1* mammary gland eQTL (P_eQTL_ = 2.64 E-12) (Liu et al., 2022b). Both eQTL lead variants were also highly significantly associated with early lactation MUC in our GWAS (P_corrected_ = 1.7 E-18 and P_corrected_ = 1.1E-18), only slightly less significant than rs431919726. The presence of closely linked regulatory and coding variants associated with MUC supports earlier suggestions that multiple alleles at the *DGAT1* locus influence dairy traits in cattle (Kühn et al., 2004).

To investigate if the association of the *DGAT1* QTL with MUC and yield traits in HOL is due to pleiotropic effects or a vertical relationship, we applied a Mendelian randomization approach under different exposure–outcome specifications (see Methods). The Mendelian randomization analysis between MUC and MKG did not yield a significant association when MKG was the exposure, but it found a significant association when MUC was the exposure (Supplemental

Table S3; see Notes), suggesting MUC-related effects on MKG. The analysis between MUC and FKG was significant in both exposure-outcome directions. This, together with a relatively low genome-wide genetic correlation of 0.15 between both traits (Supplemental Figure S2; see Notes), supports a partially shared genetic basis for MUC and FKG. After excluding instruments from the BTA14 QTL region encompassing *DGAT1*, the MUC ↔ FKG GSMR evidence was markedly attenuated compared with the genome-wide analysis. This pattern is compatible with pleiotropic variants at the *DGAT1* locus contributing to the detectable MUC–FKG GSMR signal.

*DGAT1* expression is markedly upregulated during negative energy balance (NEB) (Wang et al., 2024) that commonly occurs at lactation onset in dairy cows (McArt et al., 2013). *DGAT1* is involved in hepatic fatty acid storage and metabolic regulation (Villanueva et al., 2009). Altered hepatic *DGAT1* activity can enhance amino acid catabolism, thereby increasing blood urea concentrations (Weiner et al., 2015) which may eventually lead to higher MUC and contribute to the early-lactation specificity of this QTL.

### Evidence for multiple independent MUC-associated QTL at BTA6

A QTL for MUC on BTA6 (lead SNP: 6:85601140; P_corrected_ = 2.7E-19) was detected across all lactation stages in HOL, but the strongest association signal occurred in the late lactation stage. The lead variant was within the cluster of casein genes that harbor QTL for dairy traits (Pegolo et al., 2018; Van Den Berg et al., 2022; Atashi et al., 2024). Conditional and joint multiple-SNP analysis (COJO) suggested that this region potentially contains multiple independent QTL that impact MUC distinctly across the lactation cycle (Supplemental Table S2; see Notes). The analysis conditioned on the lead SNP provided evidence for a distant independent QTL which was observed across all windows but had the strongest association for MUC in the late lactation (lead SNP: 6:75209285; P_corrected_ = 1.54E-12). The conditional analysis also detected a QTL with effects on MUC in the mid lactation with the lead SNP residing at 85,521,678 bp (P_corrected_ = 8.5E-09) and a QTL with the lead SNP residing at 86,157,535 bp (P_corrected_ = 2.8E-09) with effects on MUC in mid and late lactation, where the association was strongest in late lactation. The genomic region encompassing the casein cluster was only suggestively associated with MUC in BSW in the mid lactation (lead SNP: 6:85612347; P_corrected_ = 6.7E-08). The top variant for the suggestive BSW QTL had a very low allele frequency in HOL (MAF =0.002), whereas the lead SNPs for the HOL QTL had high p-values in BSW. This suggests substantial heterogeneity at this locus across dairy breeds.

### Two QTL for MUC reside on BTA9 and BTA28

The GWAS identified two QTL on BTA9 (∼21.39 Mb) and BTA28 (∼6.52 Mb) in BSW that, to the best of our knowledge, have yet to be reported to be associated with MUC. Both QTL were associated with MUC in early and mid-lactation, while the association with MUC in the late lactation was weaker (BTA9) or absent (BTA28). There were different lead variants across the lactation stages, but they were all in high LD for both the BTA9 and BTA28 QTL. We inspected 1 Mb regions centered on the lead SNPs from the early lactation stage GWAS in BSW to identify candidate variants and genes (Figure 2).

**Figure 2:**
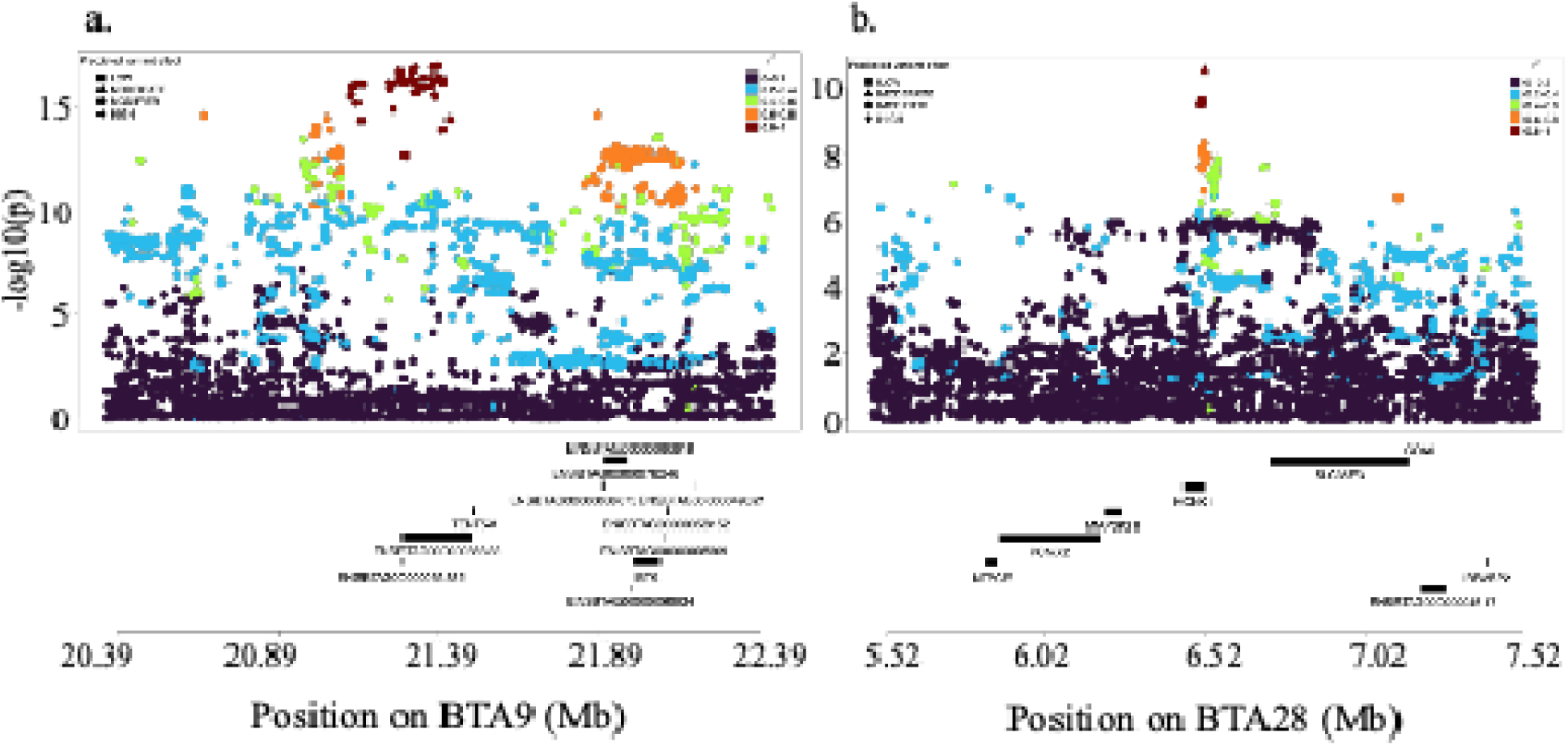
Fine mapping of two QTL for MUC in BSW cattle. Zoom plot of a 2 Mb window centered on the lead SNP (9:21392941) for a BTA9 QTL (a) and the lead SNP (28:6518357, rs133101552) for a BTA28 QTL. SNPs are colored according to their linkage disequilibrium with the lead variant. The shape of the symbol represents the VEP-predicted functional consequence of a variant. The gene content within the 2 Mb window is visualized below the zoom plot.

The lead variant for the early lactation MUC QTL on BTA9 (9:21392941; P=1.1E-17) was annotated as a non-coding variant with a putative modifier effect on the lncRNA ENSBTAG00000058688. None of the SNPs that exceeded the genome-wide significance threshold at this QTL were in coding regions (Figure 2a). ENSBTAG00000058688 contained several putative transcription factor-binding motifs (TF), including one candidate binding site for *NR1I3* (Supplemental Table S4; see Notes). *NR113* is a key metabolic regulator (Konno et al., 2008), whose role in feeding efficiency has been reported in Nellore cattle (Alexandre et al., 2014). ENSBTAG00000058688 also contains multiple annotated RNA-editing sites (Cai et al., 2025), suggesting regulatory potential. However, since the regulatory function of ENSBTAG00000058688 remains unknown, further investigation is needed to establish a potential functional link with dairy traits.

The *IBTK* gene encoding inhibitor of bruton tyrosine kinase is also a plausible functional and positional candidate gene at the BTA9 QTL although it is more distant to the lead variant than ENSBTAG00000058688. Multiple intragenic *IBTK* variants exceed the genome-wide significance threshold and were partially tagged by the lead SNP (Figure 2a). *IBTK* is highly conserved across mammals (Spatuzza et al., 2008; Dyer et al., 2025), and expressed in multiple tissues (Benoit et al., 2024; Qusairy and Rada, 2025). The cattle FarmGTEx identified an *IBTK* liver eQTL with a lead eVariant at 9:22085801(P_eQTL_ = 1.4E-12). This eVariant was strongly associated with MUC in BSW (strongest association in mid lactation, P_corrected_= 2.5E-14) and in moderate LD (r² = 0.60) with the lead variant at the BTA9 QTL. Reduced IBTK activity may enhance BTK-mediated inflammation, increasing amino-acid catabolism, and urea production (Clark et al., 1977; Zheng et al., 2023; Benoit et al., 2024; Tavakoli et al., 2024). This could possibly explain an association of this QTL primarily in the early lactation stage where inflammatory stress and mastitis susceptibility are high in dairy cattle (Zhang et al., 2024)

### Missense variant in *KCNK1* at BTA28 prioritized as a candidate

All variants exceeding the significance threshold at the BTA28 QTL overlap the *KCNK1* gene encoding the potassium two pore domain channel subfamily K member 1 (Figure 2b). The most significantly associated variant (chr28:6518357C>A, rs133101552, P_corrected_=3.0E-11) was annotated as a missense variant (KCNK1: p.Gln312Lys) with a SIFT score of 1, indicating that it is tolerated. *KCNK1* encodes a two-pore domain K channel that contributes to background K conductance and affects membrane potential in epithelial cells (Shima et al., 2024; Dyer et al., 2025).

The p.Gln312Lys-variant (rs133101552) of *KCNK1* has not previously been associated with MUC. However, it was identified as the lead variant for Fourier-transform mid-infrared spectroscopy (FT-MIR) wavenumber phenotypes in milk from New Zealand dairy cattle, which were predominantly crossbreds of Holstein-Friesian and other breeds (Tiplady et al., 2021). The same QTL region was also associated with milk MIR spectra in Holstein cows (Fresco et al., 2025) and lactose percentage in milk of Fleckvieh, Holstein, Jersey, and crossbred cows, supporting that this QTL influences milk composition (Benedet et al., 2019; Costa et al., 2019). Although rs133101552 was not significantly associated with MUC in our HOL GWAS cohort, its effect direction was consistent with that observed in BSW. Although direct evidence is lacking, *KCNK1* could affect MUC via K□-channel–driven changes in osmolality that induce the urea transporter *SLC14A1* (Alberts B; Johnson A; Lewis J; Raff M; Roberts K; Walter P, 2002; Zacchia et al., 2016; Zhu et al., 2018; Farrell and Stewart, 2019; Prahl et al., 2023; Reyer et al., 2024; Shima et al., 2024). As a canonical progesterone-responsive mammary gene, its expression may be highly sensitive to hormonal fluctuations in lactation cycle, which could help explain the stage-specific association signal (Richer et al., 2002; Gray et al., 2022; Hannan et al., 2023; Hou et al., 2024).

## Conclusions

We mapped temporal QTL dynamics for MUC across lactation windows in two Swiss dairy cattle breeds, showing that stage-specific MUC signals capture meaningful biology. Using these dynamics, we refined pleiotropy at BTA14, clarified the complex architecture at BTA6, and prioritized candidate genes for two QTL on BTA9 and BTA28 linked to immune regulation and osmotic homeostasis.

### Notes

This study was financially supported by the Arbeitsgemeinschaft Schweizerischer Rinderzüchter (ASR), Zollikofen, Switzerland, and the Federal Office for Agriculture (FOAG), Bern, Switzerland. The authors gratefully acknowledge Swissherdbook, Braunvieh Schweiz, and Holstein Switzerland for providing access to the pedigree, phenotype, and genotype data used in this study. The genotype and phenotype data analyzed in this study are owned by the respective breeding organizations and are not publicly available. Full summary statistics from the GWAS of all milk traits are available from the corresponding author upon reasonable request.

Supplemental material for this article is available at Supplement.docx. The authors declare no conflicts of interest.

