## Supplementary Material for "The genetic architecture of milk urea concentration in dairy cattle differs across the lactation cycle"

Supplemental Figure S1. Daily heritability estimates ($h^2$) for milk urea concentration (MUC) across days in milk (DIM), derived from posterior means of covariance components from the RRTDM for routine monitoring. Boxplots summarize $h^2$ estimates of $h^2$ for MUC from DIM 5 to DIM 305 in first, second, and third lactation in HOL (panels a-c) and BSW (panels d-f). Only every fifth DIM is shown. In all daily $h^2$ curves, the trajectory showed four distinct segments across DIM, corresponding to the four residual variance classes defined in the model to account for residual heterogeneity.

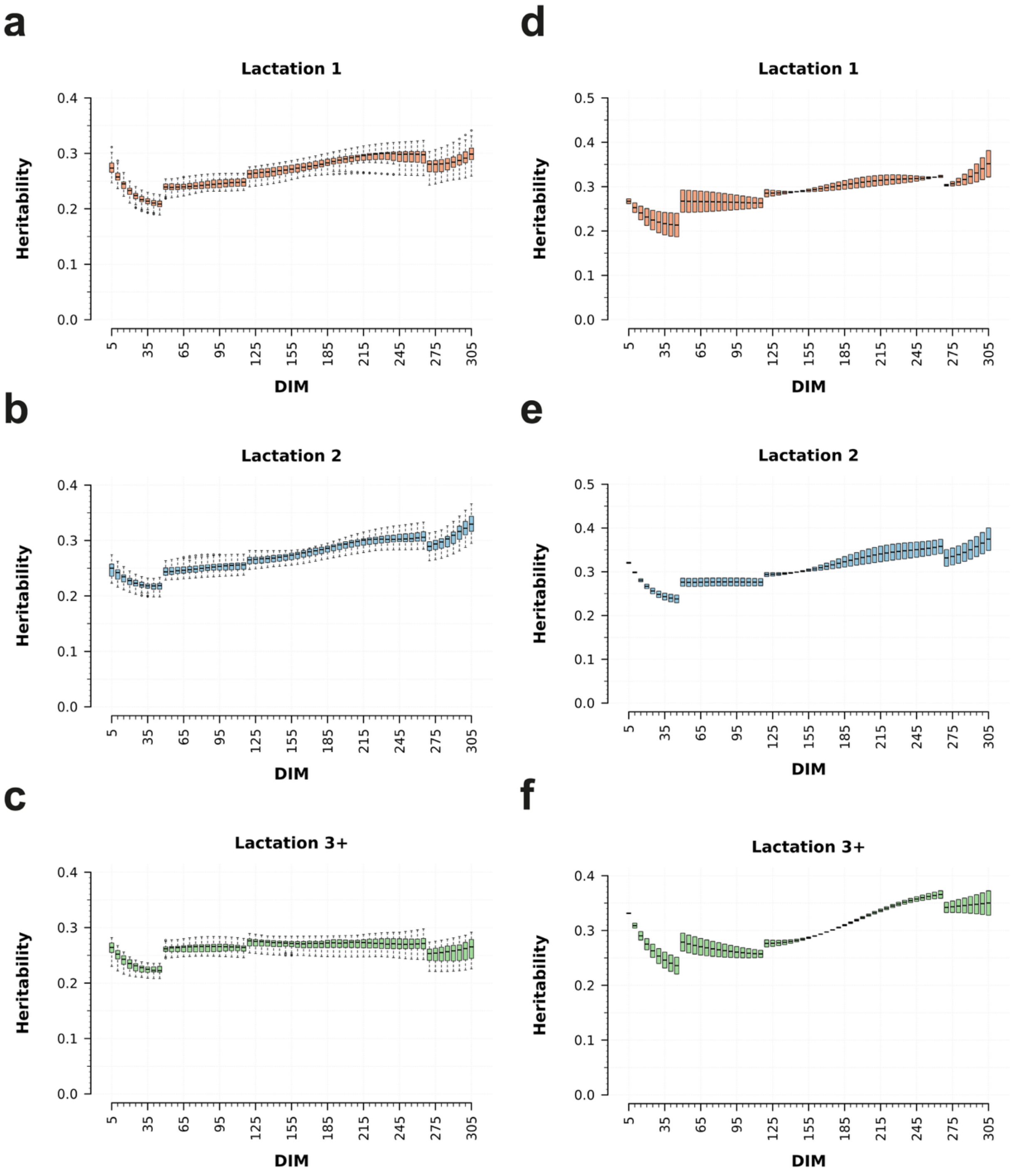
a
Lactation 1
Heritability
DIM
b
Lactation 2
Heritability
DIM
c
Lactation 3+
Heritability
DIM
d
Lactation 1
Heritability
DIM
e
Lactation 2
Heritability
DIM
f
Lactation 3+
Heritability
DIM
5
35
65
95
125
155
185
215
245
275
305

Supplemental Table S1. Milk Urea phenotype summary. The file contains the phenotype of MUC residual (Yield Deviation) from the RRTDM model that removes fix effect. For each lactation, the mean and standard deviation are also calculated.

| lactation stages | breed | mean | sd |
|---|---|---|---|
| early | BSW | 4.298 | 5,174 |
| mid | BSW | 7.014 | 4.293 |
| late | BSW | 7.322 | 4.538 |
| early | HOL | 6.032 | 4.817 |
| mid | HOL | 8.092 | 4.407 |
| late | HOL | 7.869 | 4.623 |

Supplemental Figure S2. Genetic correlation between milk urea concentration (MUC), milk yield (MKG), fat yield (FKG), and protein yield (PKG) across three lactation stages in BSW and HOL. Genetic correlations between trait pairs were estimated with GCTA. Red and blue colour represent negative and positive correlations, respectively. Values for HOL and BSW are in the upper and lower diagonal, respectively.

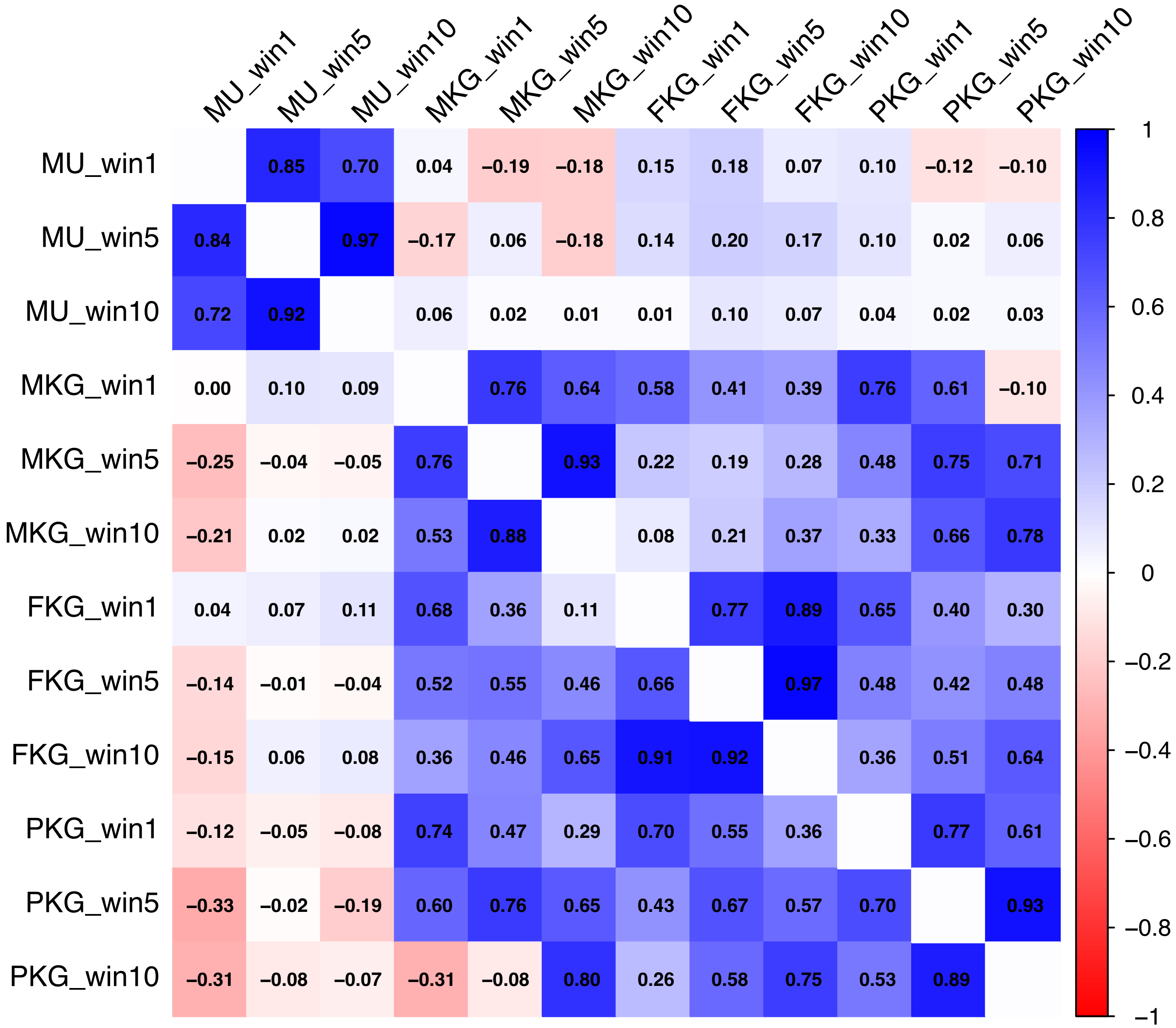
MU_win1 MU_win5 MU_win10 MKG_win1 MKG_win5 MKG_win10 FKG_win1 FKG_win5 FKG_win10 PKG_win1 PKG_win5 PKG_win10
MU_win1 0.85 0.70 0.04 −0.19 −0.18 0.15 0.18 0.07 0.10 −0.12 −0.10
MU_win5 0.84 0.97 −0.17 0.06 −0.18 0.14 0.20 0.17 0.10 0.02 0.06
MU_win10 0.72 0.92 0.06 0.02 0.01 0.01 0.10 0.07 0.04 0.02 0.03
MKG_win1 0.00 0.10 0.09 0.76 0.64 0.58 0.41 0.39 0.76 0.61 −0.10
MKG_win5 −0.25 −0.04 −0.05 0.76 0.93 0.22 0.19 0.28 0.48 0.75 0.71
MKG_win10 −0.21 0.02 0.02 0.53 0.88 0.08 0.21 0.37 0.33 0.66 0.78
FKG_win1 0.04 0.07 0.11 0.68 0.36 0.11 0.77 0.89 0.65 0.40 0.30
FKG_win5 −0.14 −0.01 −0.04 0.52 0.55 0.46 0.66 0.97 0.48 0.42 0.48
FKG_win10 −0.15 0.06 0.08 0.36 0.46 0.65 0.91 0.92 0.36 0.51 0.64
PKG_win1 −0.12 −0.05 −0.08 0.74 0.47 0.29 0.70 0.55 0.36 0.77 0.61
PKG_win5 −0.33 −0.02 −0.19 0.60 0.76 0.65 0.43 0.67 0.57 0.70 0.93
PKG_win10 −0.31 −0.08 −0.07 −0.31 −0.08 0.80 0.26 0.58 0.75 0.53 0.89
1
0.8
0.6
0.4
0.2
0
−0.2
−0.4
−0.6
−0.8
−1

Supplemental Table S2. Lead variants at MUC QTL detected in two breeds across three lactation stages. Summary statistics of lead variants at independent QTL from GWAS of MUC in BSW and HOL cattle across three lactation stages.

| CHR | POS | Breeds | lactation | freq | BETA | SE | Pgwas | Pgwas_corrected | pcojo | VEP |
|---|---|---|---|---|---|---|---|---|---|---|
| 1 | 146904166 | HOL | early | 0.4 | -0.404 | 0.0560 | 1.30E-12 | 4.80E-08 | 1.36E-12 | intergenic_variant |
| 6 | 75209838 | HOL | early | 0.12 | -0.64 | 0.085 | 2.5E-14 | 4.6E-09 | 1.7E-22 | intergenic_variant |
| 6 | 85600522 | HOL | early | 0.2 | 0.698038 | 0.071 | 1.1E-22 | 4.7E-14 | 4.65923E-66 | intronic_variant |
| 11 | 103247445 | HOL | early | 0.45 | -0.44715 | 0.058 | 8.7E-15 | 2.4E-09 | 9.56983E-15 | intergenic_variant |
| 14 | 598800 | HOL | early | 0.43 | 0.672927 | 0.058 | 5.4E-31 | 5.5E-19 | 8.60781E-31 | 3_prime_UTR_variant |
| 19 | 41767230 | HOL | early | 0.309 | -0.432823 | 0.06 | 6.6E-13 | 0.000000032 | 7.06E-13 | intergenic_variant |
| 9 | 21392941 | BSW | early | 0.31 | 0.651 | 0.055 | 4.3E-32 | 1.107E-17 | 6.85E-32 | intron_variant&non_coding_ |

| | | | | | | | | | | |
|---|---|---|---|---|---|---|---|---|---|---|
| | | | | | | | | | | transcript_variant |
| 11 | 103251967 | BSW | early | 0.33 | -0.5 | 0.057 | 1.1E-17 | 4.9E-10 | 1.2E-17 | upstream_gene_variant |
| 19 | 43963326 | BSW | early | 0.43 | 0.47 | 0.052 | 3E-19 | 7.46E-11 | 3.56E-19 | intergenic_variant |
| 28 | 6518357 | BSW | early | 0.36 | -0.502 | 0.055 | 5.5E-20 | 3E-11 | 6.56E-20 | missense variant |
| 1 | 146904166 | HOL | mid | 0.41 | -0.55 | 0.055 | 1.2E-23 | 1.5E-14 | 4.6E-20 | intergenic_variant |
| 6 | 75209838 | HOL | mid | 0.12 | -0.6 | 0.08 | 4.8E-13 | 0.000000029 | 3.97516E-12 | intergenic_variant |
| 6 | 85521678 | HOL | mid | 0.04 | -1.02 | 0.136 | 5.8E-14 | 8.5E-09 | 1.23945E-13 | intergenic_variant |
| 6 | 85866947 | HOL | mid | 0.12 | 0.855 | 0.083 | 4.2E-25 | 2.1E-15 | 5.2458E-17 | intergenic_variant |
| 6 | 86157535 | HOL | mid | 0.1 | -0.47 | 0.066 | 4.7E-13 | 0.000000029 | 6.4712E-10 | NA |
| 11 | 103254079 | HOL | mid | 0.47 | -0.51 | 0.055 | 3.1E-20 | 1.6E-12 | 3.736E-20 | upstream_gene_variant |

| | | | | | | | | | | |
|---|---|---|---|---|---|---|---|---|---|---|
| 6 | 85612347 | BSW | mid | 0.07 | 0.68 | 0.09 | 4.3E-14 | 0.000000067 | 1.63359E-12 | intergenic_variant |
| 8 | 103499852 | BSW | mid | 0.13 | -0.53 | 0.068 | 6.8E-15 | 2.58E-08 | NA | intron_variant |
| 9 | 21066396 | BSW | mid | 0.38 | -0.548 | 0.046 | 3.5E-33 | 9.4E-18 | 5.69E-33 | intergenic_variant |
| 11 | 103269281 | BSW | mid | 0.39 | -0.53 | 0.049 | 1.4E-27 | 1.3E-14 | 6.5E-27 | downstream_gene_variant |
| 19 | 45834647 | BSW | mid | 0.28 | -0.401 | 0.05 | 8.8E-16 | 8.9E-09 | 9.7E-16 | intron_variant |
| 28 | 6547458 | BSW | mid | 0.36 | -0.37 | 0.046 | 1.1E-15 | 9.8E-09 | 1.17E-15 | intergenic_variant |
| 1 | 146847984 | HOL | late | 0.42 | 0.492 | 0.067 | 1.9E-13 | 1.8E-09 | 2.55E-13 | intergenic_variant |
| 6 | 75209285 | HOL | late | 0.13 | -0.84 | 0.1 | 6.6E-18 | 1.5E-12 | 1.63379E-57 | downstream_gene_variant |
| 6 | 85601140 | HOL | late | 0.18 | 0.96 | 0.087 | 1E-28 | 2.7E-19 | 6.66475E-62 | intron_variant |
| 6 | 86882049 | HOL | late | 0.29 | -0.52 | 0.075 | 2E-12 | 0.000000013 | 2.01424E-10 | intergenic_variant |

| | | | | | | | | | | |
|---|---|---|---|---|---|---|---|---|---|---|
| 7 | 42773712 | HOL | late | 0.5 | 0.482 | 0.8 | 5.8E-12 | 0.000000016 | NA | intergenic_variant |
| 9 | 21384623 | BSW | late | 0.348 | -0.41 | 0.06 | 5.8E-12 | 0.000000041 | 6.1E-12 | intron_variant&non_coding_transcript_variant |

Supplemental Figure S3. Results of lactation stage-specific GWAS for three dairy traits in BSW and HOL cattle. Manhattan plots representing the association of imputed sequence variants with milk yield (MKG), fat yield (FKG) and protein yield (PKG) in early (a), mid (b) and late (c) lactation in BSW (left) and HOL (right). The most significant association detected across the three traits is shown for each SNP. The red horizontal line marks the genome-wide significance threshold (−log10(5E-08)). SNPs exceeding the significance threshold are colored by the trait (green = PKG, blue = MKG, red = FKG).

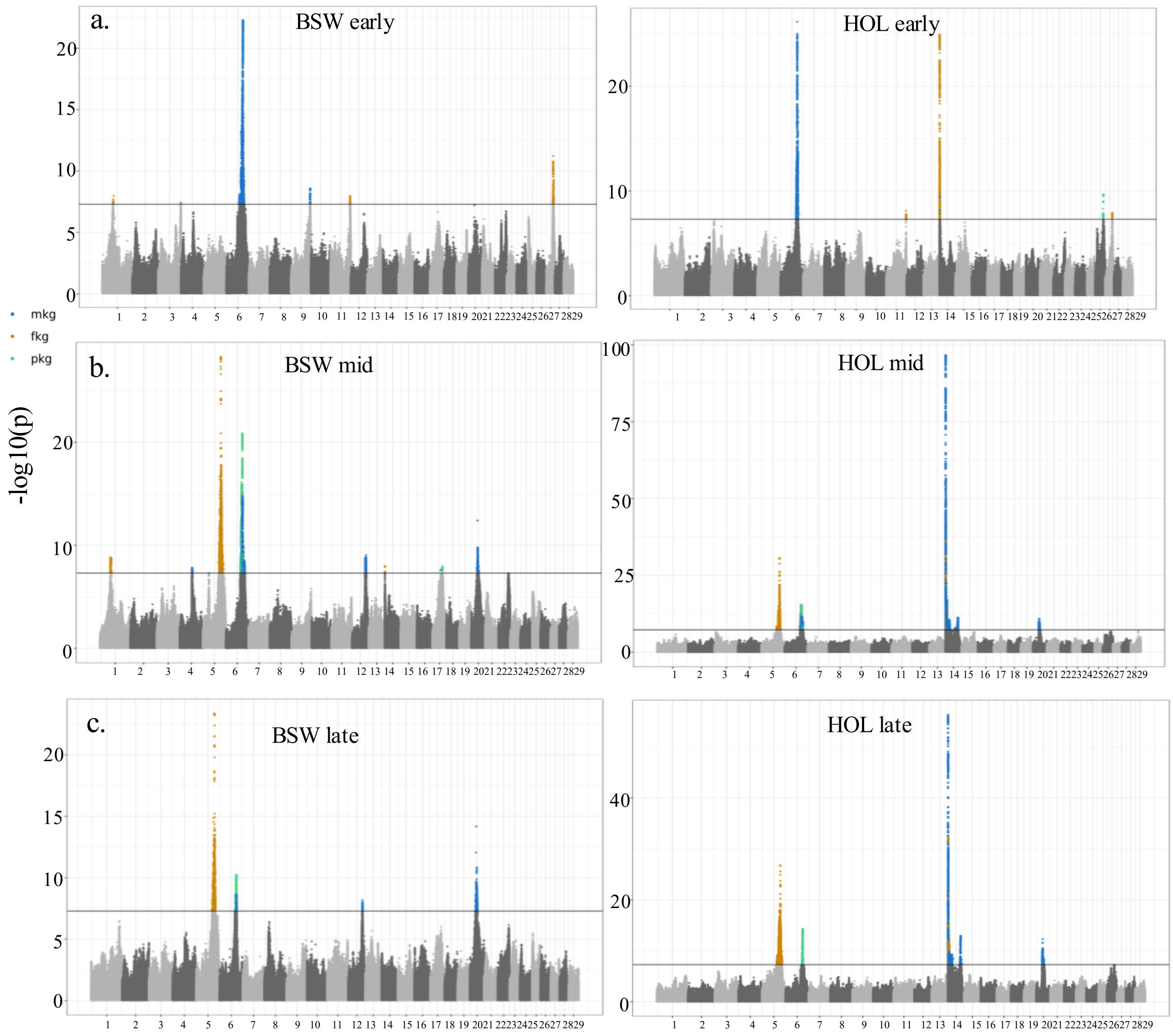

a.
BSW early
HOL early
mkg
fkg
pkg
b.
BSW mid
HOL mid
-log10(p)
c.
BSW late
HOL late

Supplemental Table S3. Mendelian randomization results for MKG, FKG and MUC. Results of the Mendelian randomization analysis with different exposure-outcome combinations for genome-wide markers, and for an analysis excluding the BTA14 region.

| Exporsure | outcome | type | effect | Pvalue | P threshold (0.05/IVs) | significant |
|---|---|---|---|---|---|---|
| MUC | MKG | whole-genome | -0.458 | 3.66E-158 | 0.00060241 | Y |
| MUC | FKG | whole-genome | 0.024 | 7.94E-98 | 0.00106383 | Y |
| FKG | MUC | whole-genome | 25.47 | 1.00E-118 | 0.00227273 | Y |
| MKG | MUC | whole-genome | -0.06 | 0.04 | 0.00217391 | N |
| MUC | MKG | BTA14 region excluded | -0.05 | 6.15E-05 | 0.00038462 | Y |
| MUC | FKG | BTA14 region excluded | -0.0021 | 1.45E-03 | 0.00054945 | N |
| FKG | MUC | BTA14 region excluded | 0.657 | 6.00E-02 | 0.0003876 | N |
| MKG | MUC | BTA14 region excluded | 0.017 | 1.00E-02 | 0.0001497 | N |

Supplemental Table S4. Candidate transcription factor (TF) binding motifs identified by FIMO. Motif occurrences were scanned using FIMO against TF position weight matrices from the Cis-BP database. The table reports the most significant predicted TF binding site for each genomic region, together with the corresponding genomic coordinates, strand, match score, and multiple-testing–adjusted significance (q value).

| MOTIF | Target gene | chr | Start | end | Strand | Score | p | q | Match length |
|---|---|---|---|---|---|---|---|---|---|
| M08377_2.00 | ZSCAN16 | 9 | 21369626 | 21369650 | - | 34.9 | 1.03E-13 | 1.60E-08 | 11 |
| M04806_2.00 | SPIC | 9 | 21283587 | 21283600 | - | 19.4 | 1.27E-08 | 0.0377 | 6 |
| M08274_2.00 | ENSBTAG00000030348 | 9 | 21386532 | 21386559 | - | 26.1 | 4.22E-10 | 0.00196 | 13 |
| M08305_2.00 | ZNF436 | 9 | 21406727 | 21406742 | + | 23.5 | 1.03E-08 | 0.0481 | 7 |
| M10879_2.00 | IRF2 | 9 | 21412950 | 21412963 | + | 20.4 | 1.44E-07 | 0.0121 | 6 |
| M09242_2.00 | IRF2 | 9 | 21460095 | 21460115 | - | 26.6 | 1.82E-12 | 7.17E-06 | 9 |
| M09315_2.00 | NR1I3 | 9 | 21428697 | 21428713 | + | 20.8 | 4.38E-09 | 0.0204 | 7 |
| M07926_2.00 | BCL11B | 9 | 21464409 | 21464424 | + | 20.3 | 1.65E-08 | 0.0239 | 7 |

| | | | | | | | | | |
|---|---|---|---|---|---|---|---|---|---|
| M07296_2.00 | LIN54 | 9 | 21438902 | 21438917 | + | 19.5 | 1.26E-08 | 0.0315 | 7 |
| M10892_2.00 | IRF2 | 9 | 21412950 | 21412963 | + | 21.7 | 1.10E-07 | 0.00581 | 6 |
| M09233_2.00 | IRF2 | 9 | 21460095 | 21460115 | - | 25.9 | 1.82E-12 | 7.90E-06 | 9 |
| M07298_2.00 | LIN54 | 9 | 21438901 | 21438916 | + | 22.1 | 3.80E-09 | 0.0151 | 7 |
